# FroM Superstring to Indexing: a space-efficient index for unconstrained *k*-mer sets using the Masked Burrows-Wheeler Transform (MBWT)

**DOI:** 10.1101/2024.10.30.621029

**Authors:** Ondřej Sladký, Pavel Veselý, Karel Břinda

## Abstract

The growing volumes and heterogeneity of genomic data call for scalable and versatile *k*-mer-set indexes. However, state-of-the-art indexes such as Spectral Burrows-Wheeler Transform (SBWT) and SSHash depend on long non-branching paths in de Bruijn graphs, which limits their efficiency for small *k*, sampled data, or high-diversity settings. Here, we introduce FMSI, a superstring-based index for arbitrary *k*-mer sets that supports efficient membership and compressed dictionary queries with strong theoretical guarantees. FMSI builds on recent advances in *k*-mer superstrings and uses the Masked Burrows-Wheeler Transform (MBWT), a novel extension of the classical BWT that incorporates position masking. Across a range of *k* values and dataset types – including genomic, pangenomic, and metagenomic – FMSI consistently achieves superior query space efficiency, using up to 2–3× less memory than state-of-the-art methods, while maintaining competitive query times. Only a space-optimized version of SBWT can match the FMSI’s footprint in some cases, but then FMSI is 2–3× faster. Our results establish superstring-based indexing as a robust, scalable, and versatile framework for arbitrary *k*-mer sets across diverse bioinformatics applications.

## Introduction

The exponential growth of DNA sequencing data calls for efficient approaches for their compression and search [36, 57]. To address the growing complexity of data, modern bioinformatics methods increasingly rely on *k*-mer tokenization for data storage and analysis. This strategy enables a unified representation of various types of genomic data, including sequencing reads, transcripts, assemblies, or species pangenomes. Notable applications of *k*-mer-based methods include large-scale data search [5, 2, 10, 28], metagenomic classification [58, 11], rapid diagnostics [6, 9], and transcript abundance quantification [7, 44]. An essential low-level functionality is the storage, and membership and dictionary queries on single *k*-mer sets [38, 37].

A key challenge is to design *k*-mer set data structures that can scale with the set size and simultaneously adapt to the structural characteristics of different sets. Comprehensive databases now reach hundreds of billions of *k*-mers [28] and continue to grow rapidly. At the same time, their structural complexity is increasing due to denser sampling of genetic polymorphisms and the accumulation of technological artifacts. Although advances in *k*-mer sampling [58] and sketching [43] techniques sets can alleviate size issues in some applications, the resulting *k*-mer sets are then less amenable to standard compression techniques.

The efficiency of *k*-mer set data structures is tightly connected to the compressive properties of their support representations. Real-world *k*-mer sets are largely non-random, and a wisely chosen indexable representation allows one to exploit redundancies, going beyond the traditional information-theoretic lower bounds [17]. Of particular benefit has been the so-called *spectrum-like property*, stating that *k*-mer sets are typically generated by a small number of long strings [13]. As these strings correspond to paths in the associated de Bruijn graphs (dBG), indexing dBG paths enables efficient indexes for the original *k*-mer sets.

State-of-the-art approaches to indexing dBG paths typically use one of two approaches: either they proceed implicitly by indexing successor *k*-mers in the graph [4, 1], or they enumerate a path cover of the dBG explicitly and then index it using either hash-based [39, 45] or full-text [14, 47, 8, 39, 45] techniques. We refer to such explicit path-cover representations as *Spectrum-Preserving String Sets* (SPSS), and refer to the literature for more details [8, 47, 51, 52]. Efficient SPSS computation is now supported by a wide range of software tools [15, 41, 26, 8, 47, 51, 52, 31, 30, 29, 18].

However, the compression power of dBG-based methods is fundamentally limited by the existence of (*k* − 1)-long overlaps between the indexed *k*-mers. Even when a fraction of *k*-mers do not share (*k* − 1)-long overlaps with other *k*-mers, long paths in the graph cannot be formed, and the corresponding *k*-mer pairs cannot be jointly compressed, impacting all these indexing methods. The resulting fragmentation causes time and space inefficiencies at all levels, especially in applications involving complex metagenomic or pangenomic *k*-mer sets, or when the sets are subsampled.

*Masked superstrings* [55] eliminate the dependency on (*k* − 1)-long overlaps. A masked superstring consists of a superstring of the original *k*-mer set, along with an associated mask that encodes the introduced “false positive” *k*-mers. Masked superstrings can represent arbitrary *k*-mer sets in text form while leveraging overlaps of all possible lengths. Conceptually, this corresponds to replacing dBGs with overlap graphs to construct the path cover. Masked superstrings thus unify all SPSS-based representations while enabling further data reduction by using shorter *k*-mer overlaps as well. Importantly, approximately shortest masked superstrings can be efficiently computed using greedy approaches implemented in KmerCamel [55] (https://github.com/ondrejsladky/kmercamel), starting from either the original *k*-mer set or a precomputed SPSS.

However, masked superstrings are not indexable using existing methods. While the superstring itself can be indexed with full-text indexing, this yields an inexact data structure with false positives. A straightforward inclusion of masks would require recovering the positions of *k*-mers in the support superstring, which in turn necessitates auxiliary data structures, such as the subsampled suffix array, thereby increasing both the space and time requirements of queries [8, 50]. Furthermore, efficient query processing in real-world applications often relies on streaming consecutive *k*-mers; yet, masked superstrings are incompatible with existing BWT-based streaming techniques [50, 49, 1] due to complications arising from *k*-mer canonicity when minimizing index size. Finally, in downstream applications requiring index associativity, to retain high space efficiency even for subsampled *k*-mer sets, the index must support the full functionality of compressed *k*-mer dictionaries [45].

Here, we introduce FMSI, a scalable data structure and software for space-efficient membership and dictionary queries over *k*-mer sets with arbitrary structure, with support for accelerated streamed queries. FMSI combines compact *k*-mer set representation using approximately shortest masked superstrings [55] with indexing via the Burrows-Wheeler Transform (BWT) [12], extended to accommodate for masks. The resulting index provides strong theoretical guarantees on space for arbitrary *k*-mer sets, including those resulting from sampling and sketching. We implement this in a program called FMSI (https://github.com/ondrejsladky/fmsi) and demonstrate its superior space efficiency across a broad range of datasets and use cases compared to current state-of-the-art *k*-mer-set indexing methods.

## Methods

### Problem formulation

Let us consider the nucleotide alphabet {*A, C, G, T* }. For a fixed *k* > 0, *k-mers* are strings of length *k* over this alphabet. We assume the *bidirectional model* (formalized, e.g., in [15, online suppl.],[51]), where a *k*-mer and its reverse complement (RC) are considered equal, unless the *unidirectional model* is explicitly indicated. We are interested in providing the following indexing functionality for a *k*-mer set *K*.

(P1) **Membership**

⊳ Single *O*(*k*)-time operation: ⊳ Member(*Q*): Determine if a given *k*-mer *Q* is in *K*.

(P2) **Dictionary**

Two *O*(*k*)-time operations enabling a bijective mapping between *k*-mers *Q* ∈ *K* and a discrete interval of hash values *H* = {0, …, |*K*| − 1}:

⊳ Lookup (*Q*): Return the hash *h* ∈ *H* of *Q*, or −1 if *Q* ∉ *K*.
⊳ Access (*h*): Return the *k*-mer *Q* corresponding to *h* ∈ *H*.

(P3) **Streaming**

Member/Lookup in *O*(1) time for a *k*-mer *Q* following a query for *k*-mer *Q*^′^ when the (*k*−1)-long prefix of *Q* is a suffix of *Q*^′^ and both *Q, Q*^′^ ∈ *K*.

The dictionary index (P2) corresponds to evaluating a *minimal perfect hash function* with rejection and its inverse. (P2) is more general than (P1), but (P1) can be implemented more efficiently. (P3) corresponds to extending the core data structure for isolated queries by a component for rapid fwd or bwd operations (as defined in [13]), thus trading a small additional space for improving the time efficiency of positive streamed queries.

Our goal is to develop a data structure that simultaneously addresses four challenges. First, it should minimize space requirements while maintaining competitive query speed. Second, it must make no assumptions about the structure in the data, such as about the presence of many (*k* − 1)-long overlaps. Third, when structural regularities such as overlaps are available, the method should be able to exploit them. Fourth, the data structure should be available in four optimized variants, each targeting a specific functionality: a membership index (P1), a compressed dictionary (P2), and two extended versions with streaming support (P1+P3 and P2+P3, respectively).

### Two core ingredients of FMSI

Our approach is based on indexing approximately shortest superstrings of the input *k*-mers, using a technique based on the BWT [12] (**Fig. 1**). Depending on the desired functionality (**Fig. 1a**), we compute a particular variant of the masked superstring representation [55], obtain the BWT image of superstring together with an appropriate mask transformation, and complete the data structure with supporting rank and select data structures into an exact index termed FMSI (**Fig. 1b**). FMSI shares similarities but is different from existing BWT-based approaches such as dbgFM [14, 47], BWA [34, 8]/ProPhex [49, 50], and SBWT [1] (**Fig. 1c, Tab. 1**).

**Table 1.** Functional comparison of FMSI with state-of-the-art *k*-mer indexes. Comparison of the supported operations for single-set *k*-mer indexes, including FMSI and state-of-the-art methods.

|  | P1: Memb. | P2: Dict. | P3: Stream. | Scalable | Dynamic |
| --- | --- | --- | --- | --- | --- |
| BWA [34] | ✓ [8] | ✗ | ProPhex [49, 50] | ✓ | ✗ |
| dbgFM [14, 47] | ✓ | ABYSS [54] | ✗ | ✗ | ✗ |
| SSHash [45] | ✓ | ✓ | ✓ | ✓ | ✗ |
| CBL [40] | ✓ | ✗ | ✓ | ✓ | ✓ |
| SBWT [1] | ✓ | ✗ | ✓ | ✓ | ✗ |
| FMSI | ✓ | ✓ | ✓ | ✓ | [56] |

**Fig. 1:**
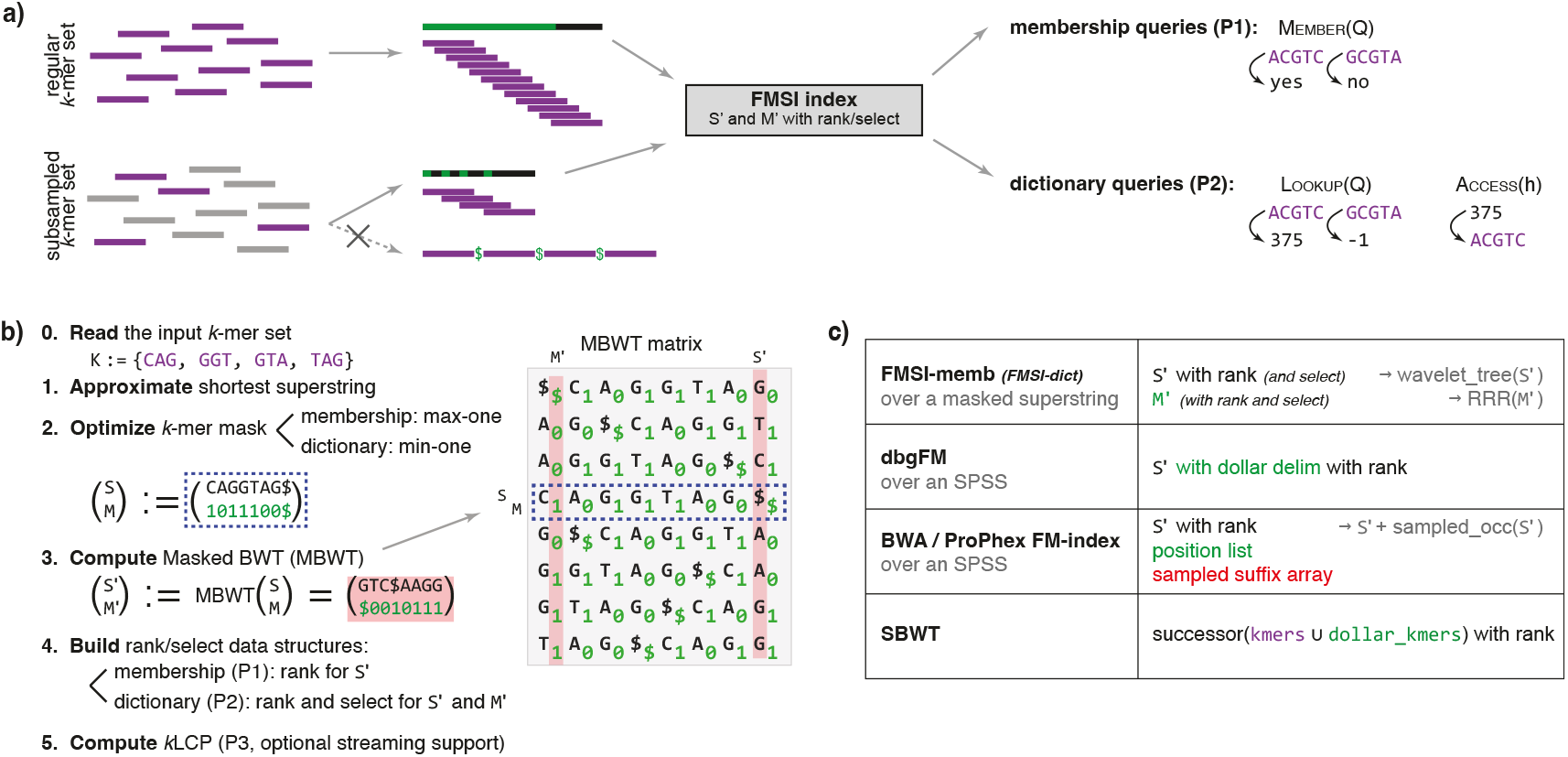
Overview of the FMSI method. Concepts related to input *k*-mers and masks are shown in violet and green, respectively. **a) Core FMSI functionality**. FMSI indexes regular or subsampled *k*-mer sets and supports both membership or dictionary queries. Unlike other approaches, its representation remains compact even without (*k* − 1)-long overlaps. **b) FMSI index construction**. A greedy global algorithm [55] approximates the shortest superstring of the input *k*-mer set; the resulting mask is optimized for the target functionality (P1 or P2). The masked superstring is transformed via the Masked BWT (**Def**. **1** and **Alg**. **1**) and equipped with rank/select data structures. Optionally, the *k*LCP array [49] can be computed to accelerate streamed queries. **c) Comparison with other BWT-based indexes**. dbgFM [14, 47] indexes $-separated SPSS using an FM index without a sampled inverse suffix array. BWA/ProPhex [34, 8, 49, 50] index SPSS concatenations, avoiding false matches at simplitig borders by checking simplitigs’ start positions. SBWT [1] indexes dBG paths via *k*-mer successors. Only FMSI removes redundancy among *k*-mers with overlaps shorter than *k* − 1.

#### 1. Approximately shortest masked superstrings of *k*-mers

A *masked superstring* representation [55] of a given *k*-mer set *K* consists of two parts: a superstring *S* of the *k*-mers, i.e., a single string containing all the *k*-mers in *K* as substrings, and a binary mask *M* determining which of the *k*-mers appearing in *S* are represented. A *k*-mer is considered represented by a masked superstring (*S, M*) if at least one of its occurrences in *S* has the corresponding mask symbol set to 1, and the set of represented *k*-mers should be equal to *K*. Since it is sufficient to mask each *k*-mer as present only once, this makes the mask not unique, allowing for additional mask optimization. Masked superstrings unify all SPSS representations and provide a more general framework with better compression properties [55].

Here, we consider uniquely masked superstrings obtained via greedily approximated shortest superstrings [55] and, based on the target functionality, masked to have either the maximum or the minimum number of ones, termed max-one or min-one masked superstrings, respectively (Steps 1, 2 in **Fig**. **1b**). Given input *k*-mer data, such a superstring can be efficiently approximated by the global greedy algorithm as implemented in KmerCamel [55], with possible acceleration via parallelized SPSS precomputation. The output min-one masked superstring can further be converted to max-one using a two-pass algorithm [55].

#### 2. Masked Burrows-Wheeler Transform

We index the resulting masked superstring using the *Masked Burrows Wheeler Transform*, an extension of the traditional BWT [12] with support for masks in the suffix array (SA) order (**Fig**. **1b**).

##### Definition 1

(MBWT) Let (*S, M*) be a masked superstring and let *S*^′^ be the BWT image of *S*. The *SA-transformed mask* is a string *M*^′^ of the same length as *M* such that for all *i, M*^′^[*i*] = *M*[*j*] where *j* is the starting position of the *i*-th lexicographically smallest suffix of *S*. We call the pair (*S*^′^, *M*^′^) the *Masked Burrows-Wheeler Transform* (MBWT) of (*S, M*).

The SA-transformed mask can be obtained alongside the standard BWT via a small tweak: we start by rotating the mask by one position to the left, “glue” the mask symbols to the superstring symbols, and then compute the BWT of the superstring with respect to the lexicographical order based solely on the superstring characters (i.e., disregarding the mask bits in the string comparisons). We formalize the whole process in **Alg. 1** and prove its correctness in **Lemma 1**.

##### Lemma 1

(SA-transformed mask construction) *Let* (*S, M*) *be a masked superstring, M*^′^ *the SA-transformed mask, S*^′^ *the BWT of S, and* 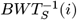 *the original coordinates in S of the i-th character from S*^′^.*Then* 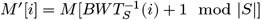.

*Proof* Consider the *i*-th lexicographically smallest rotation of *S*. The last character of this rotation has position 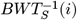 in *S* and since the first character cyclically succeeds it in *S*, it has coordinate 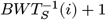 mod |*S*| in *S*. Therefore, *M*^′^[*i*] must be 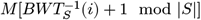.

##### Algorithm 1

MBWT computation.

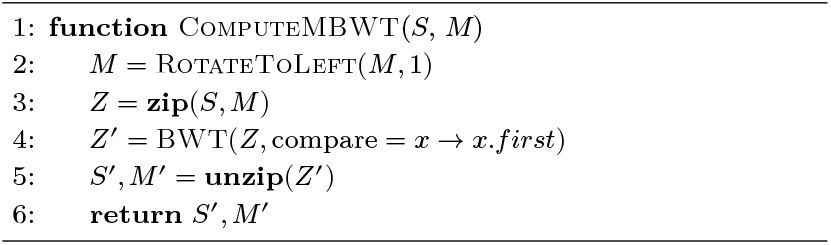

##### Algorithm 2

*k*-mer search in FMSI

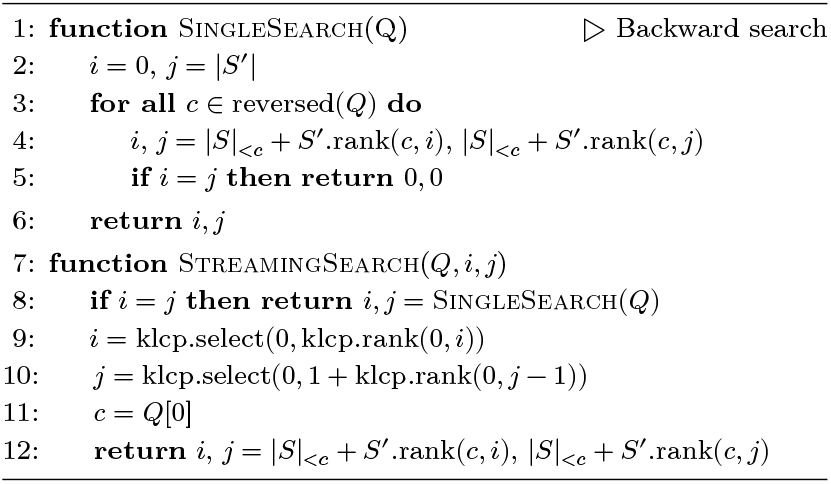

##### Algorithm 3

Query operations (given SA range [*i, j*) or hash *h*)

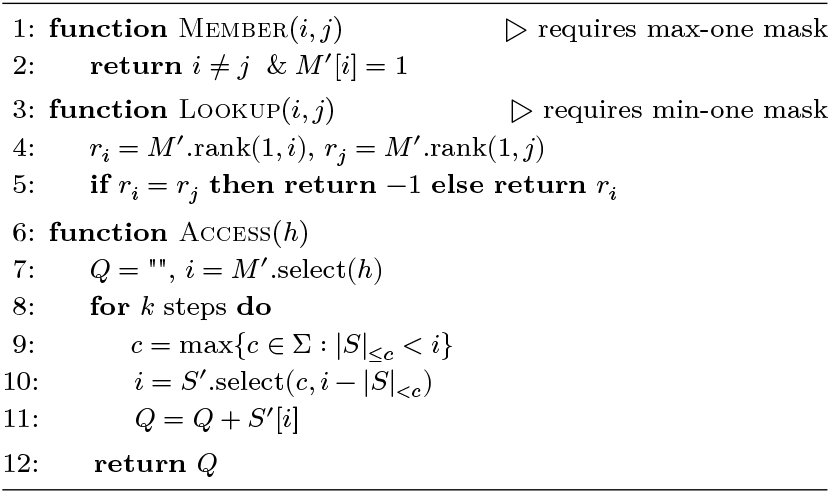

Therefore, MBWT can be constructed with any method for BWT computation that allows alphabets with eight symbols, to represent all pairs (*s, m*) of superstring and mask symbols, while considering pairs (*s, m*) and 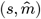 with the same superstring symbol as lexicographically equivalent even if 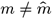, that is, using only *s* for comparisons.

### The FMSI membership and dictionary data structures

The FMSI data structure combines the two ingredients with suitable supporting data structures. FMSI indexes an approximately shortest masked superstring (*S, M*) via its MBWT image (*S*^′^, *M*^′^), which is complemented by the rank data structure [27] for *S*^′^. In the case of the dictionary variant of FMSI, it additionally requires select [16] for *S*^′^ and rank and select for *M*^′^ (**Fig**. **1b**).

The memory footprint of FMSI is tightly connected with the (*S*^′^, *M*^′^) representation, whose size should be minimized. Since approximately shortest *k*-mer superstrings are typically not compressible beyond two bits per *k*-mer, a natural representation for *S*^′^ is a plain bit-vector. In contrast, the masks tend to be highly imbalanced: towards all-zero vectors for subsampled *k*-mer sets, and towards all-one vectors with the spectrum-like property [55].

Therefore, *S*^′^ can be represented using rank and select succinct data structures [27, 16], and *M*^′^ can be encoded via zeroth-order entropy compression in data structures such as the *succinct indexable dictionary* [48, 22], with support for constant-time rank and select operations.

#### Definition 2

(FMSI) Given a *k*-mer set *K*, a corresponding masked superstring (*S, M*), and its associated MBWT image (*M*^′^, *S*^′^), we define the following data structures:

- **FMSI-memb** = (*S*^′^ with rank, *M*^′^), assuming that *M maximizes* the number of 1’s in the mask, for membership queries (P1).
- **FMSI-dict** = (*S*^′^ with rank and select, *M*^′^ with rank and select), assuming that *M minimizes* the number of 1’s in the mask, for dictionary queries (P2).

### FMSI index theoretical properties

We now analyze the theoretical guarantees of the FMSI index construction and its space requirements. First, we analyze index construction time, starting from a suitable (max-one or min-one) masked superstring (*S, M*) of the indexed *k*-mer set *K*.

#### Lemma 2

*Given a masked superstring* (*S, M*) *of a k-mer set K, both FMSI-memb and FMSI-dict data structures can be constructed in expected time O*(|*S*|).

*Proof* The MBWT image (*S*^′^, *M*^′^) can be computed in linear time with respect to the superstring length (**Alg. 1**). All aforementioned data structures for rank and select can be constructed in *O*(|*S*|) time as well; only for the compressed representation of *M*^′^ using the succinct indexable dictionary, the construction runs in *expected* linear time [22]. □

Second, we analyze the storage requirement of FMSI – first, in the general case of arbitrary *k*-mer sets, and then when the underlying superstring size approaches the number of distinct canonical *k*-mers (e.g., only 4% difference for the human genome and *k* = 31; **Tab. 3**).

#### Lemma 3

(FMSI storage requirements) *Given a masked superstring* (*S, M*) *of a k-mer set K, the corresponding FMSI-memb and FMSI-dict indexes require* 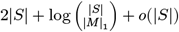 *bits, which is at most* (3 + *o*(1)) · |*S*|. *Furthermore, if* |*S*| = |*K*| + *o*(|*K*|), *the index requires* 2 + *o*(1) *bits per distinct canonical k-mer*.

*Proof S*^′^ stored as a plain bit-vector-based wavelet tree [24] requires 2 + *o*(1) bits per character, including the rank and select support. *M*^′^ stored using a zeroth-order entropy compression [48, 22] requires 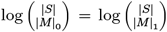 bits. Both rank and select for the compressed vector *M*^′^ occupy only sublinear space [48, 22]. Last, suppose that |*S*| = |*K*| + Δ with Δ ∈ *o*(|*K*|). Letting 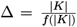 for *f*(|*K*|) → ∞ as |*K*| → ∞, the space complexity of *M*^′^ is 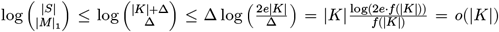 and, thus, the index requires 2 + *o*(1) bits per distinct *k*-mer. □

### FMSI membership and dictionary queries

FMSI can act as two slightly different data structures, a membership index and a dictionary, with membership queries being slightly more space- and time-efficient (**Tab. 2**). In both cases, the search starts with the standard backward search, as used, e.g., in the FM index, resulting in a suffix array (SA) interval corresponding to occurrences of the query *k*-mer in the superstrings (**Alg**. **2**). The backward search uses the “count” values for characters *c*, denoted |*S*|_<*c*_ and equal to the total number of occurrences of all characters *c*^′^ < *c* in *S*, which can be computed during the FMSI construction in constant space. The candidate *k*-mer matches in the SA intervals, if non-empty, are subsequently validated against *M*^′^ to exclude false positives or to obtain the *k*-mer’s hash (**Alg**. **3**). In the bidirectional model, everything works the same way with only one difference: If the forward *k*-mer is not found in the superstring, we subsequently query also the RC of the *k*-mer.

**Table 2.** Query time and memory usage for membership vs. dictionary queries. Comparison of query time (per query *k*-mer, *μ*s) and memory (bits per distinct canonical *k*-mer, in *italics*) on three datasets with *k* = 31. See Experimental Evaluation for details.

| Dataset | FMSI-memb | FMSI-dict |
| --- | --- | --- |
| ec-pg-hq | 17.2 \| 3.3 | 20.5 \| 3.4 |
| mtg-ilm | 7.7 \| 3.6 | 14.6 \| 3.7 |
| hg-t2t | 11.4 \| 2.4 | 14.8 \| 2.4 |

**Table 3.** Index construction resources on human genome (*k* = 31). CPU time of construction (usr+sys, minutes) and memory usage (GB, in *italics*) for the human genome dataset. The greedily computed superstring had length |*S*| = 2.602 G, with |*K*| = 2.512 G. See Experimental Evaluation for details (the **hg-t2t** dataset).

| Method | SPSS | MS | Index | Overall |
| --- | --- | --- | --- | --- |
| CBL | — | — | 38.4 \| 101.1 | 38.4 \| 101.1 |
| SBWT-small | — | — | 21.5* \| 49.9 | 21.5* \| 49.9 |
| SBWT-fast | — | — | 39.5* \| 93.1 | 39.5* \| 93.1 |
| SSHash<sup>†</sup> | 30.5* \| 21.6 | — | 14.6 \| 10.9 | 45.1 \| 21.6 |
| FMSI-memb | 30.5* \| 21.6 | 25.3 \| 39.0 | 15.4 \| 44.2 | 71.2 \| 44.2 |
| FMSI-dict<sup>†</sup> | 30.5* \| 21.6 | 14.1 \| 1.7 | 18.8 \| 44.2 | 63.4 \| 44.2 |

<sup>†</sup> indexes supporting dictionary queries \* with 8 threads (1 otherwise)

### Membership queries

In FMSI-memb, Member is evaluated by inspecting the first bit of the SA-transformed mask within the interval [*i, j*) obtained via backward search (**Alg**. **3**). This is possible thanks to the max-one property of the mask, which ensures that all bits in the interval are set to the same value.

### Dictionary queries

FMSI-dict supports the Lookup operation via additional rank data structure, required for two additional invocations of *M*^′^.rank to compute *r*_*i*_ = *M*^′^.rank(1, *i*) and *r*_*j*_ = *M*^′^.rank(1, *j*) (**Alg**. **3**). If *r*_*i*_ = *r*_*j*_, the *k*-mer is not in *K* and is reported as absent from the dictionary; otherwise, *r*_*i*_ is returned as its unique minimal hash. This is possible thanks to the min-one property of the mask, which ensures that *r*_*i*_ equals the number of distinct represented *k*-mers appearing before the queried *k*-mer in the SA order, making *r*_*i*_ both unique and minimal.

In the other direction, Access requires select data structures for both *S*^′^ and *M*^′^. It is evaluated via *M*^′^.select(*h*), which returns the position *i* of the *h*-th occurrence of 1 in *M*^′^. This position marks the start of the SA interval corresponding to the queried *k*-mer *Q*, which we then reconstruct in *k* steps by iteratively applying select on *S*^′^ (**Alg**. **3**).

### Queries with unoptimized masks

Membership operations are possible with masks other than max-one, but at the cost of additional time overhead due to two invocations of rank on *M*^′^ to determine the presence of 1 in the SA interval via *M*^′^.rank(1, *i*) < *M*^′^.rank(1, *j*). Likewise, dictionary operations are possible with masks other than min-one, but at the cost of losing the contiguity and surjectivity of hash values, as their range length increases by |*M*|_1_ − |*K*|.

### Streaming acceleration

In many practical scenarios, *k*-mers are queried in batches consisting of all overlapping *k*-mers of a given query string, such as a sequencing read or a genomic element. Oftentimes, the streamed queries are almost entirely positive, e.g., when a query gene differs only in one nucleotide due to a single mutation. By *positive streamed queries*, we mean a query for (all occurrences of *k*-mers of) a string *T* such that all the *k*-mers of *T* are present in the represented *k*-mer set.

To accelerate positive streamed queries, we use the *k*LCP array technique, originally developed in the context of *k*-mer indexing with FM-indexes [49] and previously implemented in ProphEx [50]. This technique is based on the bit-vector of the same length as *S* such that the *i*-th bit is set to 1 if and only if the longest common prefix between the *i*-th and (*i* + 1)-th suffixes of *S* in the lexicographic order has length at least *k* − 1. This allows to obtain a range corresponding to a (*k*−1)-mer obtained after deleting the last character. In the unidirectional model, this gives us immediately fast positive streamed queries, in the bidirectional model, we need to ensure that we query most *k*-mers first in the direction in which they appear in the superstring.

### Unidirectional streaming via *k*LCP

Without taking RCs into account, we traverse the query sequence in the reverse direction (**Alg. 2**). Starting from the last *k*-mer, after we have queried a *present k*-mer, we obtain the range for its prefix of size *k* − 1 by extending the range in the SA coordinates in both directions to the first 0 in the *k*LCP array using two rank and select operations. To get the range for the next *k*-mer, we perform an update of the range with its first character, i.e., one step of the backward search, which only costs *O*(1) time in total. Only when the current queried *k*-mer does not appear in the superstring, we perform the backward search for the next *k*-mer in time *O*(*k*).

### Streaming bidirectionality via saturating strand counters

In the bidirectional model, consecutive queried *k*-mers may change directionality in the superstring; however, the *k*LCP extension technique supports only one fixed direction. Nevertheless, streaming can still be accelerated locally if we can split queries into locally consistent parts and query them in the right direction. As a simple but effective heuristic, we split queries into blocks of size *B*, and for each block we estimate the likely dominating directionality of its *k*-mers via a saturating counter. Specifically, whenever a *k*-mer is found on the forward strand, we increment the counter, and decrement otherwise (and vice versa for the reverse strand). Then, the directionality of the next block can be estimated based on the statistics stored in this counter, informing whether the block should be first reverse-complemented or not.

This technique provides favorable theoretical time guarantees for positive streamed queries, both in the unidirectional and bidirectional model (**Lemma 4**). For the unidirectional model we have slightly better guarantees, whose difference, however, vanishes as the size of the query sequence grows. The speedup in the bidirectional model assumes that consecutive *k*-mers do not change strand in the superstring too often; for example, this readily holds for greedy superstrings built from *k*-mer sets corresponding to de Bruijn graphs with limited branching.

#### Lemma 4

*Given an FMSI index over a masked superstring* (*S, M*) *of a k-mer set K, and a query sequence T containing t* = |*T* | − *k* + 1 *occurrences of k-mers from K, streamed queries can be processed such that:*

- *In the unidirectional model, querying T takes time O*(*t* + *k*) = *O*(|*T* |), *or* 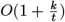 *per k-mer*.
- *In the bidirectional model, if there are at most O* (*t*/*k*^2^) *pairs of consecutive k-mers Q*_*i*_, *Q*_*i*+1_ *in T, such that Q*_*i*+1_ *does not appear on the same strand as Q*_*i*_ *in S, then querying k-mers in T takes time* 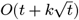, *or* 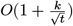 *per k-mer*.

*Proof sketch*. Querying the last *k*-mer of *T* takes *O*(*k*) time. In the unidirectional model, since all queries are positive, every other *k*-mer of *T* can be queried in *O*(1) time as explained above.

In the bidirectional model, let 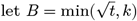. We split *T* into blocks of size Θ(*B*). For each of these blocks, we predict whether the *k*-mer or its RC are in *S* using the saturating counter with small constant saturation computed on previous blocks, which predicts to start with the strand with which the previous block ended with. We process the first block in time at most *O*(*B* · *k*). For other blocks, we spend time *O*(*k*) on the first *k*-mer. Then, if we predict the strand correctly and the strand does not change during the block, we spend time *O*(*k* + *B*) per block, and otherwise, we spend time *O*(*k*) for each *k*-mer. The total time spend on blocks in the former case is thus 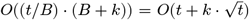. The latter can happen if consecutive *k*-mers switch strands either inside the block, in the previous block, or between blocks which may cause an incorrect prediction. However, such switching happens at most *O* (*t*/*k*^2^) times in total and hence, the total time complexity is bounded by 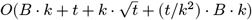, which is always bounded by 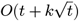. □

### Implementation in the FMSI program

We implemented the FMSI index in a tool called FMSI, an abbreviation of ‘FroM Superstring to Indexing’. FMSI is developed in C++, provided on Github under the MIT licence (https://github.com/OndrejSladky/fmsi), deposited on Zenodo (https://doi.org/10.5281/zenodo.13905020), and distributed via Bioconda [25]. The version of FMSI used in this paper is 0.4.0. FMSI can calculate an index for user-provided masked superstring (*S, M*) for an arbitrary *k*, and then query it with Membership or Lookup queries, possibly streamed. The input masked superstring can be computed by KmerCamel (https://github.com/OndrejSladky/kmercamel), or alternatively inferred from any SPSS representation.

The index in the FMSI software consists of three components: *S*^′^ as a BWT image of *S* (stored via a wavelet tree over non-compressed plain bit-vectors), *M*^′^ as the corresponding SA-transformed mask (an RRR-compressed bit-vector [48]), and an optional *k*LCP array (a plain bit-vector), all based on the sdsl-lite library [23]. MBWT computation proceeds via suffix array construction using QSufSort [32].

For single-*k*-mer queries, we also utilize the strand prediction technique we described for streamed queries. If a *k*-mer is not found in the superstring (**Alg. 2** and **3**), its RC is also queried, starting from a predicted strand (using a counter with saturation 7). FMSI uses two additional counters, to accumulate and predict only if the previous *k*-mer was found in the forward or reverse direction respectively. Streaming (**Alg. 2**) additionally simplifies the range extension: the *k*LCP array is traversed in both directions until 0s are found, avoiding the additional rank data structures. Bidirectionality is supported via splitting each queried sequence of length *ℓ* into parts of size 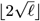. The initial query strand of each part is predicted from the saturating counters of the preceding parts, with only the first strand queried entirely and the other one only for the not-yet-found *k*-mers.

### Experimental Evaluation

#### Experimental setup

We benchmarked FMSI against three state-of-the-art indexes for single *k*-mer sets: *SBWT* (Spectral Burrows-Wheeler Transform [1], commit b795178), *SSHash* ([45], version 3.0.0), *CBL* (Conway-Bromage-Lyndon [40], commit 328bcc6), each configured as recommended by their authors. Specifically, SSHash used minimizer sized as min{⌈log _4_|*K*|⌉ + 1, *k* − 2} and indexes based on optimal simplitigs (eulertigs) computed by GGCAT [18]. CBL was independently compiled per *k*-mer size, with SIMD support and 28 prefix bits. SBWT was evaluated in two distinct modes: a standard mode optimized for speed (*SBWT-fast*, with a plain matrix and index with RCs) and mode tuned for minimum memory (*SBWT-small*, with RRR-split and index without RCs), both considered with and without streaming support. *DbgFM* [14] was excluded due to scalability issues and recurrent index construction failures.

We selected eight datasets representative of diverse tradeoffs between *k*-mer set sizes and sets’ structural properties common across bioinformatics applications: SARS-CoV-2 pangenome (**sc2-pg**; n=14M; GISAID [53]), *S. pneumoniae* pangenome (**sp-pg**; n=616; [19]), *E*.*coli* 661k pangenomes [3] with (**ec-pg-hq**; n=86K) and without (**ec-pg-all**; n=89K) quality filtering, metagenomic Illumina reads of a human tongue microbiome (**mtg-ilm**; SRX023459),

RNA-seq Illumina reads (**rna-ilm**; SRX348811), human T2T assembly (**hg-t2t**; chm13v2.0 [42]), and human WGS Illumina reads (**hg-ilm**; SRX016231). Each dataset was preprocessed with GGCAT [18] to compute an SPSS, followed by KmerCamel [55] to compute greedy masked superstrings for FMSI. Illumina raw-read datasets (\***-ilm**) were cleaned via *k*-mer filtering (*k* = 32, min-freq 2). Example summary of construction resources for **hg-t2t** and *k* = 31 are provided in **Tab. 3**. Benchmarks were performed on an AMD EPYC 7302 (3 GHz) server with 251 GB RAM and SSD storage. Comprehensive protocols and pre-processed datasets are provided in **Supplementary material**.

#### FMSI consistently achieves best space efficiency

We systematically evaluated the memory usage of FMSI, SBWT-small, SBWT-fast, SSHash, and CBL queries across dataset types (without subsampling) and various *k*-mer sizes (**Tab**. **4**). FMSI consistently showed the lowest memory footprint. Specifically, it was by 1–2 orders of magnitude more space-efficient than CBL, over 2.5× more space-efficient than SSHash, and over 3× more space-efficient than SBWT-fast. The only method competitive in space with FMSI was SBWT-small, but only on large *k*-mer sets with dBG dominated by long paths (e.g., the human genome assembly **hg-t2t** and the *E*.*coli* pangenome without contamination filtering **ec-pg-all**, both only at large *k*). Since index serialization on disk enables additional compression and omits recomputable index parts and buffers, we also evaluated on-disk index sizes (smaller values in **Tab. 4**). The results were consistent, except for **sc2-pg**, where the SBWT-small serialized index was marginally smaller than FMSI’s at large *k*.

**Table 4.** Memory and disk usage across *k*-mer sizes for membership query indexes. Reported values are in bits per canonical *k*-mer, measured across all benchmark datasets and indicated *k*-mer sizes. Space usage was measured in memory (via GNU time; larger numbers) and on disk (smaller, italicized numbers). The best lower and upper value for each dataset is indicated by asterisk (*). Due to memory constraints (>251 GiB), CBL could not be evaluated for **hg-t2t** at *k* > 39.

| Dataset | k | 31-mers | CBL |  | SSHash |  | SBWT-fast |  | SBWT-small |  | FMSI |  |
| --- | --- | --- | --- | --- | --- | --- | --- | --- | --- | --- | --- | --- |
| **sc2-pg** | 15, 17, ..., 43 | 12.7 M | 153.8–654.1 | *46.6–78.7* | 14.0–25.0 | *10.7–19.3* | 22.2–56.0 | *9.4–10.3* | 16.0–50.8 | *3.2\*–5.0* | 5.5*–9.0* | *3.4–3.7\** |
| **sp-pg** | 15, 17, ..., 43 | 9.62 M | 150.3–300.9 | *47.5–88.9* | 8.2–22.6 | *5.5–18.0* | 29.9–45.9 | *9.2–9.8* | 24.5–41.4 | *3.6–5.1* | 6.1*–7.4* | *2.8\*–3.4\** |
| **ec-pg-hq** | 15, 17, ..., 43 | 551 M | 34.3–173.0 | *20.7–80.9* | 6.6–22.7 | *6.6–22.6* | 9.0–9.7 | *8.5–8.6* | 3.4–7.8 | *3.1–6.6* | 3.1*–3.5* | *2.8\*–3.3\** |
| **ec-pg-all** | 15, 23, ..., 31 | 1.32 G | 25.5–69.0 | *18.7–48.9* | 7.6–22.0 | *7.5–21.9* | 8.8–9.2 | *8.5–8.6* | 3.1–7.8 | *2.9–7.1* | 2.9*–3.0* | *2.7\*–2.8\** |
| **mtg-ilm** | 15, 17, ..., 31 | 355 M | 37.1–65.5 | *21.0–50.9* | 7.8–22.7 | *7.7–22.6* | 9.5–10.8 | *8.6–10.1* | 3.7–7.3 | *3.1–6.1* | 3.0*–3.7* | *2.8\*–3.4\** |
| **rna-ilm** | 15, 23, ..., 31 | 686 M | 39.6–130.4 | *20.5–48.8* | 14.9–22.8 | *14.9–22.6* | 9.7–14.4 | *8.7–14.0* | 4.8–7.8 | *4.4–6.7* | 3.3*–6.2* | *3.0\*–5.9\** |
| **hg-ilm** | 15, 23, ..., 31 | 3.56 G | 25.1–341.1 | *18.6–48.5* | 10.8–20.6 | *10.8–20.6* | 9.2–10.5 | *8.5–10.4* | 3.5–8.5 | *3.4–7.8* | 2.9*–4.1* | *2.6\*–3.9\** |
| **hg-t2t** | 15, 17, ..., 43 | 2.51 G | 21.1–597.0 | *19.0–73.1* | 5.9–24.2 | *5.9–24.2* | 8.6–9.2 | *8.5–8.5* | 2.4*–8.0 | *2.3–7.3* | 2.4*–3.0* | *2.2\*–2.9\** |
| Overall |  |  | 21.1–654.1 | *18.6–88.9* | 5.9–25.0 | *5.5–24.2* | 8.6–56.0 | *8.5–14.0* | 2.4*–50.8 | *2.3–7.8* | 2.4*–9.0* | *2.2\*–5.9\** |

#### FMSI is robust to *k*

We next examined how memory usage varied with *k* (**Fig. 2**). FMSI and SBWT-fast exhibited stable memory consumption per *k*-mer, remaining within a narrow range of the minimum observed value (except **rna-ilm**). SSHash’ performance varied largely with *k*, with the best space requirements achieved for large values of *k*; otherwise, it required up to 2–4 times more space. SBWT-small remained space efficient at large *k*, but performance was degraded by a factor of 1.6–3.3 at smaller *k*. In contrast, CBL showed the opposite trend – beyond *k* > 23, its memory exceeded 40 bits per *k*-mer. We note SSHash and CBL performance may be further tuned via targeted (dataset,*k*)-specific optimization of the minimizer size and prefix bit parameters, respectively.

**Fig. 2:**
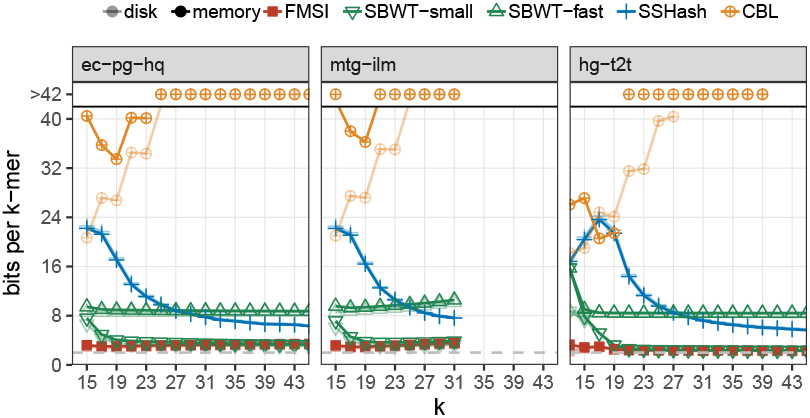
Memory usage of indexes across *k*-mer lengths. Memory usage is reported in bits per canonical *k*-mer, shown as a function of *k*. Corresponding on-disk sizes are indicated in lighter color. For **mtg-ilm**, evaluation is limited to *k* ≤ 32 due to initial *k*-mer filtering. The gray dashed line indicates two bits per *k*-mer.

#### FMSI maintains competitive query times

Next, we eval-uated whether FMSI maintains competitive query times compared to other methods (**Fig. 3a**). We selected *k* = 23 as a representative setting where all indexes achieve reasonable space usage and compared their positive and negative query times, with and without streaming acceleration. Overall, FMSI’s query speed was in the mid-range relative to the other methods. SSHash and SBWT-fast were generally the fastest, outperforming FMSI by a factor of 2–4×. At the same time, FMSI was usually 2–4× faster than the second best space-efficient tool, SBWT-small, although ties occurred, e.g., for *E. coli* pangenomic datasets with streaming support. FMSI’s use of *k*LCP for accelerated streaming improved queries 2–4× (and similarly for SBWT), but at the cost of an additional bit per superstring character. In contrast, SSHash and CBL accelerate streaming without additional memory overhead. Importantly, the relative performance of methods under streaming remained consistent with that observed in isolated query mode. Interestingly, for **hg-t2t** with accelerated streaming, FMSI achieved comparable query times as SSHash, both being approximately 2× faster than SBWT-fast.

**Fig. 3:**
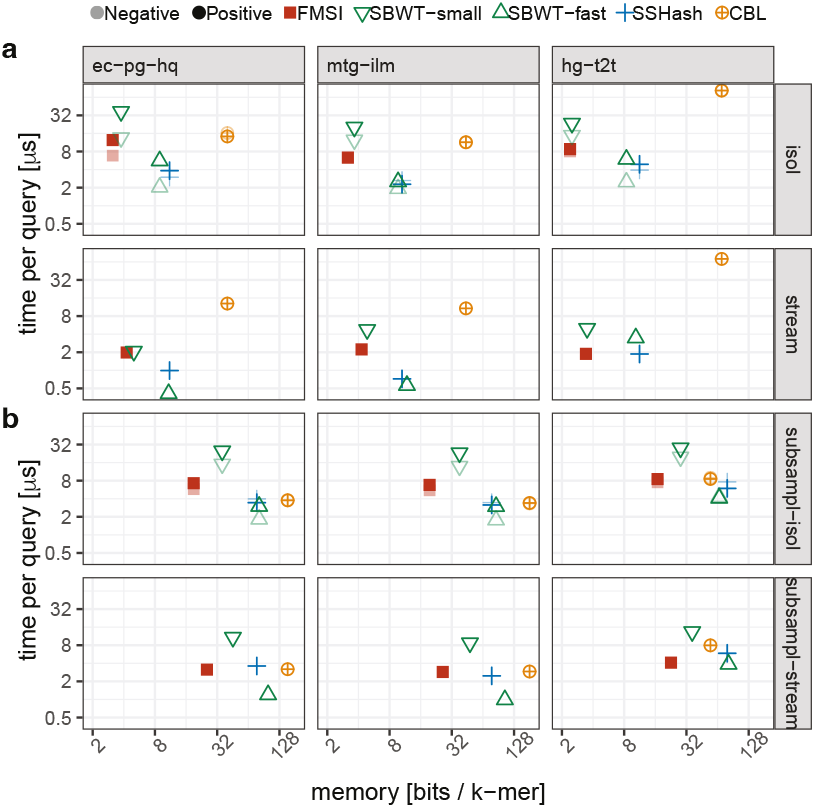
Time-memory trade-offs for membership queries for a) regular and b) subsampled *k*-mer sets (10%). Query time (per *k*-mer) and memory usage are shown for selected datasets with *k* = 23, with and without streaming acceleration, and with and without reference set subsampling.

**Fig. 4:**
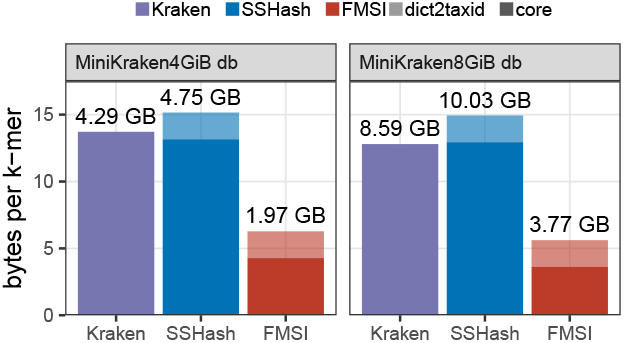
Disk usage of LCA indexes on MiniKraken database. Comparison of disk size for Kraken, FMSI, and SSHash indexes on subsampled MiniKraken v20171019 (*k* = 31). FMSI and SSHash include an additional two bytes per *k*-mer to associate each hash with the lowest common ancestor in the taxonomic tree.

#### Membership on subsampled *k*-mer sets

Next, we sought to understand the effect of sampling on the query time and space of all indexes. We uniformly subsampled the *k*-mer sets to 10% on the previous *k*-mer sets (**Fig. 3b**). This resulted in a substantial increase in memory per *k*-mer of all tested tools, except for CBL on **hg-t2t**. The memory increase is primarily due to missing overlaps of length *k* − 1 in the dataset, resulting in substantially larger representation lengths, when normalized by the *k*-mer set size. FMSI typically used at most half the space of the second best tool, SBWT-small, while also being 2× faster. FMSI was approximately 4× more space-efficient than SBWT-fast and SSHash, while being 2–3× slower on isolated queries. FMSI was particularly efficient on streamed queries, similarly fast as SSHash while using about 4× less space, achieved thanks to the “*k*-mer interpolation” nature of superstrings.

#### FMSI reduces the Minikraken index size 2×

Finally, we sought to demonstrate the utility of FMSI as a dictionary of sampled *k*-mer sets, and evaluated its compressive properties on the subsampled database of the Kraken metagenomic classifier [58] termed the MiniKraken databases (**Fig**. **4**). We built FMSI-dict and SSHash 31-mer dictionaries for both MiniKraken 31-mer databases available online (with an additional two bytes per *k*-mer for taxid arrays), and compared them to the size of the original indexes. We found that while SSHash moderately increased memory requirements compared to the Kraken index, FMSI provided more than a twofold memory improvement in both cases.

## Discussion and Conclusions

*k*-mer-based methods are central to modern bioinformatics, and efficient *k*-mer set indexing is therefore a critical objective. In this work, we introduced FMSI, a data structure and its implementation for space-efficient indexing of single *k*-mer sets. FMSI consistently achieved lower space usage than all other state-of-the-art methods across a broad operational range, while maintaining competitive query times. This was possible thanks to approximately shortest superstrings and BWT indexing – two techniques that had not previously been combined in this context.

Several performance tradeoffs in FMSI could be further improved. First, index construction currently depends on QSufSort [32], which limits its scalability, whereas modern BWT construction algorithms already scale to terabyte-sized inputs [35]. Second, query speed would benefit from parallelization, implemented similarly to BWT-based indexes [34, 1]. Third, space usage may be further reduced by applying Elias-Fano [20, 21] to the SA-transformed mask instead of RRR [48]. Finally, FMSI could be re-optimized for speed instead of space – for instance, by adding a reverse complement superstring to the index or replacing wavelet trees with 2-bit encoding, in analogy to the fast modes in SBWT.

Despite decades of work and a rich palette of available methods, *k*-mer indexing remains an exciting problem with substantial room for improvement. This is especially true in non-standard settings involving subsampling, sketching, or high levels of genetic polymorphisms and technological artifacts – such as those encountered in metagenomics and pangenomics. In these cases, superstrings offer an elegant means to bypass the limitations of traditional methods, representing a major weapon in the algorithmic arsenal for future generic, space-efficient, and parameter-free *k*-mer-based data structures. Their potential impact may extend well beyond bioinformatics, into domains such as indexing in large language models.

## Author contributions statement

O.S., P.V., K.B. developed the methodology, designed and conducted the experiments, analyzed the results, and wrote the manuscript. O.S. developed the software.

## Acknowledgments

This research was supported by the French National Research Agency (ANR) under Grant ANR-24-CE45-1226 for the REALL project (KB), by Czech Science Foundation project 22-22997S (OS, PV), by ERC-CZ project LL2406 of the Ministry of Education of Czech Republic (PV), by Center for Foundations of Modern Computer Science (Charles Univ. project UNCE 24/SCI/008, PV), and by Campus France under PHC BARRANDE 2025 grant n° 52374TC for the EFFIMAS project (KB, PV). Portions of this research were conducted at the GenOuest bioinformatics core facility (https://www.genouest.org). We are grateful to Camille Marchet, Igor Martayan, Giulio Pibiri, Jarno Alanko, and Paul Medvedev for their constructive comments.

## Supplementary material

### Experimental evaluation

All scripts, data tables, and additional information are available in the supplementary repository of our paper on https://github.com/OndrejSladky/fmsi-supplement. The experimental evaluation was done in the following five steps.

#### Step 1: Download of the source data

The *SARS-CoV-2* pangenome **sc2-pg** was downloaded from GISAID https://gisaid.org/ (access upon registration, version 2023/01/23). The *E. coli* pangenome was obtained from the phylogenically compressed 661k collection [3] as provided on https://doi.org/10.5281/zenodo.4602622 [10]; all the genomes from the batches starting by ‘`escherichia_coli`’ were extracted and used subsequently. Dataset **ec-pg-all** is the whole *E. coli* pangenome, without quality filtering, while for **ec-pg-hq** we applied high-quality filtering. The *S. pneumoniae* pangenome was downloaded from the RASE DB *S. pneumoniae* (https://github.com/c2-d2/rase-db-spneumoniae-sparc/). Metagenomic sample SRS063932 (Illumina raw reads) of human microbiome with accession SRX023459, denoted **mtg-ilm**, was download from https://www.hmpdacc.org/hmp/HMASM/; we converted the fastq files into FASTA files by ‘seqtk seq -A -C’. Human RNA-seq Illumina raw reads with accession SRX348811, **rna-ilm**, and the human genome Illumina raw reads with accession SRX016231, **hg-ilm**, were downloaded using the prefetch tool from the SRA toolkit and then converted into the FASTA format by ‘`fastq-dump --split-3 --fasta`‘. The Human genome assembly chm13v2.0, **hg-t2t**, was downloaded from https://s3-us-west-2.amazonaws.com/human-pangenomics/T2T/CHM13/assemblies/analysis_set/chm13v2.0.fa.gz. The MiniKraken datasets (4GiB and 8GiB) were downloaded from https://ccb.jhu.edu/software/kraken/

#### Step 2: *k*-mer set preparation

The input files for the pangenomes, mtg-ilm, and hg-ilm reads were converted to unitigs by GGCAT v1.1.0 [18] by `‘ggcat build -k {kmer-size} -m 200 -j 5 -s {min-freq} -o {preprocessed} {input_FASTA}’`. We used *k* = 128 and {min-freq}=1 for pangenomes and *k* = 32 and {min-freq}=2 for **mtg-ilm** and **hg-ilm**. No preprocessing was done for the HG assembly chm13v2.0 (hg-t2t). For MiniKraken datasets (4GiB and 8GiB), we dumped the 31-mers using Jellyfish 1.1.12. The obtained files with the *k*-mer sets were deposited on https://zenodo.org/records/14722244.

#### Step 3: Representation tailoring

Before indexing with SShash [45] and FMSI, we used GGCAT [18] (v1.1.1) and KmerCamel [55] (v2.1.1), respectively, for preprocessing before indexing. The following commands were used (with *k* matching the *k* passed to the programs for indexing).

- For SShash, we computed eulertigs, which is the shortest SPSS representation: ‘`ggcat build -k {kmer-size} --eulertigs --min-multiplicity 1 --memory 200 --threads-count 8 -o {eulertigs} {preprocessed}`‘.
- For FMSI with the best available MS, including the optimization to max-one mask: ‘kmercamel compute -k {kmer-size} -M {ms-maxone} -o {ms-minone} {preprocessed}’. Alternatively, MS for FMSI was computed from eulertigs: ‘kmercamel compute -S -k {kmer-size} -M {ms-maxone} -o {ms-minone} {eulertigs}‘.

#### Step 4: Generating *k*-mer queries

To generate queries, we simulated Illumina-like reads by WgSim [33] (v0.3.1-r13) with the following command:’ `wgsim -1{query_length} -d0 -S42 -s31 -e0 -r{mut_rate} -R0 -N{num_seqs} {wgsim_input} {queries} /dev/null’, where ‘query_length`’ was equal to *k* for isolated queries and to 300 for streaming queries, ‘`mut_rate`’ was 0 for positive (isolated and streaming) queries and 0.1 for negative queries, ‘`num_seqs`’ was set so that there are 10^6^ *k*-mer queries overall (up to 300 less for streaming queries), and ‘`wgsim_input`’ was dataset and query-type dependent; namely, for generating negative isolated queries, we used chromosome 1A of *T. aestivum* assembly GCF_018294505.1, downloaded from NCBI. We generated positive isolated and streaming queries from the original dataset or a reference genome in case of pangenomes (details provided in the supplementary repository). For input to the version of SBWT which does not store reverse complementary *k*-mers, we additionally duplicated every line of all query files and computed the reverse complements of the duplicated lines using ‘`seqtk seq -r`‘.

**Step 5: Construction and querying of the indexes** We describe the commands used for the construction of indexes and running the queries. Time and memory for index construction are summarized in **Tab**. **5**, including MS/SPSS computation of Step 3. The measured performance of the indexes is provided in **Tab. 2** and **Figs. 2-4**. Further information and experimental results can be found at https://github.com/OndrejSladky/fmsi-supplement.

- **FMSI** (https://github.com/OndrejSladky/fmsi/, v0.4.0, commit ‘04c4580’). FMSIhas two variants, depending on the desired query type:
  - **Variant for membership queries:** index construction by ‘`fmsi index -k {kmer-size} {ms-maxone}`’ and queries using ‘`fmsi` query `-k {kmer-size} -O -q {queries} {ms-maxone}`‘, additionally with ‘-S’ in case of streaming queries.
  - **Variant for dictionary queries:** index construction by ‘`fmsi index -k {kmer-size} {ms-maxone}`’ and lookup queries using ‘`fmsi lookup -k {kmer-size} -q {queries} {ms-minone}`‘, additionally with ‘`-S`’ in case of streaming queries.
- **SBWT** [1] (https://github.com/algbio/SBWT, commit ‘b795178’). We compiled SBWT separately for *k* ≤ 32 and for *k* ∈ (32, 64]. The specific parameters were tuned based on the results provided in [1]. As the time-efficient variant, we use the default plain-matrix variant, with adding all reverse complements to the index. As the space-efficient variant, we use the rrr-split variant without reverse complements and queried both *k*-mer and its reverse complement.
  - **Fast variant:** index construction using ‘sbwt build -k {kmer-size} -m 120 -t 8 --add-reverse-complements \\ `-i {prefix} -o {index-path}`‘, additionally with ‘`--no-streaming-support`’ for isolated queries, and queries executed by ‘`sbwt search -i {index-path} -q {queries} -o /dev/null`‘.
  - **Space-efficient variant:** index construction using ‘`sbwt build --variant rrr-split -k {kmer-size} -m 120 -t 8 -i {prefix} -o {index-path}`‘, additionally with `--no-streaming-support` for isolated queries, and queries executed by ‘`sbwt search -i {index-path} -q {queries_with_RCs} -o /dev/null`‘, always querying a *k*-mer and its reverse complement (RC).
- **SSHash** [45, 46] (https://github.com/jermp/sshash, commit ‘d90ad37’). A dictionary based on minimal perfect hashing of *k*-mers. We compiled SSHash separately for *k* ≤ 31 and for *k* ∈ [32, 63]. Indexes were computed using ‘`sshash build -k {kmer-size} -m {M} -s {seed}` ‘, where we set the minimizer length to ‘M’ = min{⌈log _4_# *k*-mers⌉ + 1, *k* − 2} and used a random seed. Whenever the index construction failed, SSHash was executed again with a different random seed.
  - **Single variant:** index construction using ‘`sshash build -i {prefix} -k {kmer-size} -m {M} -o {index-path} -s {seed}`‘, where `prefix` is the input file with an SPSS of the *k*-mer set, and queries using ‘`sshash query -i {index-path} -q {queries}`‘
- **CBL** [40] (https://github.com/imartayan/CBL, commit ‘328bcc6’), a very recent method based on smallest cyclic rotations of *k*-mers. The index was computed on canonical *k*-mers, to handle reverse complements, that is, we ran ‘`cbl build -c`‘. CBL was compiled for every value of *k* separately, namely, we used ‘`RUSTFLAGS=“-C target-cpu=native” K={kmer-size} PREFIX_BITS={pref_bits} \\ cargo +nightly build ---release ---examples ---target-dir target.k_{kmer-size}`‘ where the pref_bits were set to 28 for *k* ≥ 15, to 24 for *k* = 13, and to 20 for *k* = 11. We then used the following commands:
  - **Single variant:** index construction done using `‘cbl build -c -o {index-path} {prefix}`‘, where `prefix` is the preprocessed dataset, and queries using ‘`cbl query {index-path} {queries}`’

## Supplementary tables

**Table 5.**
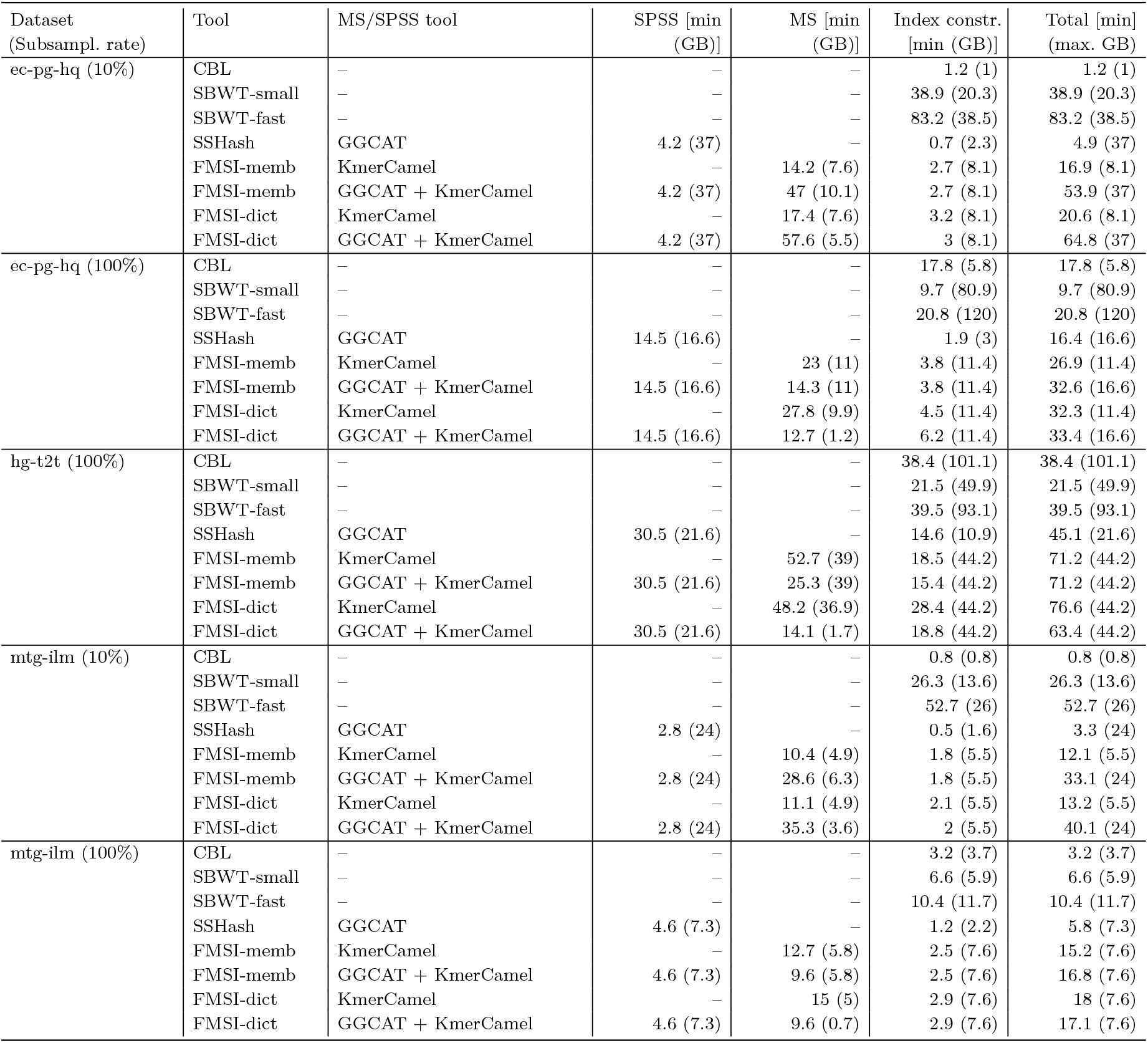
Construction time and memory usage for superstrings and indexes. Construction CPU time (usr+sys, minutes) and memory usage (GB, in parentheses) for selected datasets with *k* = 31. SPSS are optimal simplitigs (eulertigs) computed by GGCAT [18]; MS is computed using the global greedy algorithm of KmerCamel [55], either from preprocessed input or from SPSS. GGCAT and SBWT were run on 8 threads; the other tools support only single-threaded construction.

